# Histopathological Domain Adaptation with Generative Adversarial Networks Bridging the Domain Gap Between Thyroid Cancer Histopathology Datasets

**DOI:** 10.1101/2023.05.22.541691

**Authors:** William Dee, Rana Alaaeldin Ibrahim, Eirini Marouli

## Abstract

Deep learning techniques are increasingly being used to classify medical imaging data with high accuracy. Despite this, due to often limited training data, these models can lack sufficient generalizability to predict unseen test data, produced in a different domain, with comparable performance. This study focuses on thyroid histopathology image classification and investigates whether a Generative Adversarial Network (GAN), trained with just 156 patient samples, can produce high quality synthetic images to sufficiently augment training data and improve overall model generalizability. Utilizing a StyleGAN2-ADA approach, the generative network produced images with an FID score of 5.05, matching state-of-the-art GAN results in non-medical domains with comparable dataset sizes. Augmenting the training data with these GAN-generated images increased model generalizability significantly when tested on external data, improving overall accuracy and F1 scores by 36% and 42% respectively. Most importantly, this performance improvement was observed on minority class images, tumour subtypes which are known to suffer from high levels of inter-observer variability when classified by trained pathologists.

## Introduction

Thyroid cancer incidence has generally been increasing since the 1970s (1). It is currently the ninth most common cancer worldwide (2), and the 2019 Global Burden of Disease study predicted incidence will increase across all age groups for the next 20 years (3).

Differentiated thyroid cancer (DTC) includes all types of thyroid cancer that originate in the cells which produce and store thyroid hormones, and accounts for approximately 90% of thyroid cancer incidence (3, 4). DTC generally has a good prognosis compared to undifferentiated thyroid cancers which include anaplastic and medullary thyroid cancer (5). Within this DTC designation, the most common thyroid gland malignancy is papillary thyroid carcinoma (PTC), constituting 80-90% of diagnosed cases (3, 6).

PTC, along with other variants designated as PTC-like – including non-invasive follicular thyroid neoplasm with papillary-like nuclear features (NIFTP) and follicular variant of papillary thyroid carcinoma (FVPTC) – have distinctive nuclear features which aid their clinical classification (7).

These include changes to nuclear size and shape, primarily elongation, enlargement and overlapping nuclei, as well as chromatin alterations such as “clearing, margination and glassy nuclei” (8).

Whilst histopathology assessments remain the gold standard in tumour diagnosis (7, 9), there still exists significant interobserver variability between diagnoses (10, 11). Additionally, diagnostic accuracy is dependent on the experience of the pathologist (12), and greater patient imaging throughput is placing increasing demands on the time of these highly qualified professionals (13).

In the past two decades, the evolution of whole-slide image technology has facilitated the digital storage of high-resolution histopathological images. The ability to share high quality sample data globally has enabled the development of various machine learning (ML) approaches aimed at automating histopathological image classification.

Computer-based methods have the potential to improve diagnosis speed and accuracy, as they require less overall training time than a human and have been shown to outperform experienced clinicians in various image classification tasks (14, 15). Furthermore, an interpretable ML system can be used to aid pathologist training, as well as enabling quality assurance and the assessment of both inter and intra-observer variability (9).

Accurate diagnosis is particularly important within thyroid cancer as overdiagnosis is a known and growing issue, accounting for up to 60-90% of newly diagnosed cases (16, 17). This overdiagnosis places needless psychological burden on patients and can lead to overtreatment, i.e., unnecessary thyroidectomy surgery (18).

Due to PTC’s prevalence and relatively clear features, combined with long-term survival rates of more than 90% (19), identifying patient samples with PTC-like nuclei is the first step in a pathologist’s diagnostic approach (20). ML approaches have therefore often focused on automating the bulk of the diagnosis burden, classifying histopathological images as PTC-like or not (13, 20–22). These methods have utilised a variety of ML architectures, ranging from Random Forests and Support Vector Machines (SVMs) (23–25) to Convolutional Neural Network (CNN) models (20, 26, 27).

Böhland et al. (20) provided one of the first direct comparisons between six different ML methods and a deep learning CNN approach. However, the best performing models still lacked generalizability when applied to an external test set, produced in a different domain.

The authors suggested that this was likely due to a lack of diverse training data, a common problem when applying ML methods to (often small) medical imaging datasets (14). This issue can be exacerbated when using deep learning methods, which typically require large datasets to produce highly generalizable models (28). To visualize the domain gap between the training and test data, (20) passed both image datasets through a ResNet50 (29) classifier, before using Uniform Manifold Approximation and Projection (UMAP) (30), to represent the data in two dimensions.

The separation displayed in Figure 1 demonstrates that the data distributions are clearly distinct from each other – i.e., there are more differences between the datasets caused by the domain origination, than similarities between images of the same classification.

**Fig. 1.**
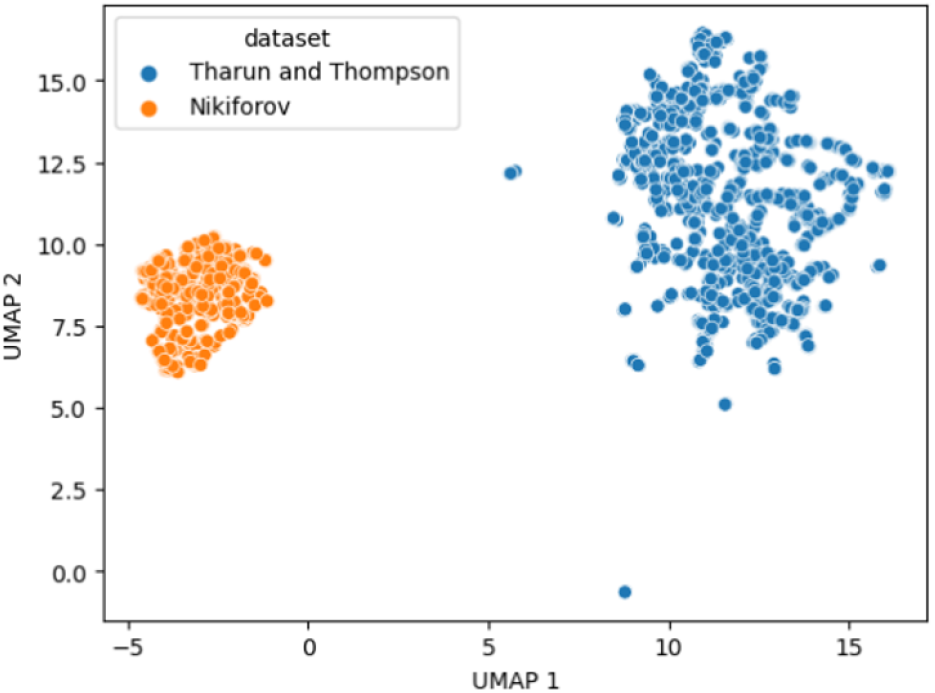
Domain gap visualization between the Tharun and Thompson dataset (20) and their external Nikiforov test data (31), using UMAP. The figure shows a clear separation between the two datasets – referred to as the “Domain Gap”. This gap can have a negative impact on the ability of a model trained on one dataset being able to generalize to predict the other. Source: Böhland et al. (20)

Generative adversarial networks (GANs) were introduced by Goodfellow et al. (32) as a method of producing high quality synthetic (fake) images which approximate an underlying real data distribution. Adversarial training methods have been applied successfully in the field of medical imaging in areas such as MRI reconstruction and tumour segmentation (33), X-ray organ segmentation (34, 35), and virtual slide staining (36–38).

GAN-generated synthetic images have been proven to pass visual Turing tests when presented to trained medical professionals. Synthetic lung nodule samples were assessed as real 67% of the time by a radiologist with 13 years’ experience, whilst 100% were considered real by a radiologist with four years’ experience (39). Both board-certified and trainee pathologists showed an inability to distinguish between real and synthetic ovarian carcinoma samples, choosing correctly only 54% of the time (40). Lastly, Xue et al. (37) found that three out of four pathologists could not differentiate over half of the synthetic histopathology images produced by their GAN.

Augmenting training data with GAN-generated images has been shown to increase the performance of ML models when generalizing to unseen test data. In the case of Liu et al. (41), the accuracy of predicting isocitrate dehydrogenase status increased 8.8% from 78.4% to 85.3%. Similarly, Guan & Loew (42) showed an uplift of 3.6% from GAN-generated mammogram images over other forms of data augmentation (i.e., affine transformations), when classifying breast cancer.

In this study we aimed to investigate whether a GAN could be successfully trained on a limited dataset of 156 patient histopathology slides to produce realistic synthetic images which could improve the generalizability of a deep learning classifier when applied to external data, produced in different domains.

## Materials and methods

### Data acquisition and processing

#### Tharun and Thompson (“T&T”) dataset

The dataset is comprised of 156 whole slide images (WSIs) of thyroid gland tumours, 138 sourced from the University Clinic Schleswig-Holstein, Campus Luebeck, and 18 from the Woodland Hills Medical Centre, California (20). Two pathologists agreed on the classification of each tumour, before 1,916 px by 1,053 px crops were extracted from the identified neoplastic regions of interest. Each image has a magnification factor of 40x and a resolution of 0.23 μm/px. The dataset was requested by emailing. See Table 1 for additional information.

**Table 1.** Sample subtypes present in the Tharun and Thompson (“T&T”) dataset, including their classification as PTC-like or not and the number of patient samples in each category.

| Subtype | Classification | Source Data | Patients |
| --- | --- | --- | --- |
| Papillary thyroid carcinoma (PTC) | PTC-like | T&T | 53 |
| Follicular variant of papillary thyroid carcinoma (FVPTC) | PTC-like | T&T | 9 |
| Non-invasive follicular thyroid neoplasm with papillary-like nuclear features (NIFTP) | PTC-like | T&T | 9 |
| Follicular thyroid adenoma (FA) | Non-PTC-like | T&T | 53 |
| Follicular thyroid carcinoma (FTC) | Non-PTC-like | T&T | 32 |
| **Total** |  |  | **156** |

PTC samples constitute the majority of the positive “PTC-like” class, whilst the FVPTC and NIFTP subtypes, which are considered much more difficult to diagnose due to their less distinctive nuclear features (20), are minority samples within the data – mimicking their real-life comparative scarcity. The “Non-PTC-like” class consists of the two most common diagnoses which lack PTC-like nuclei – FA and FTC. Additional detail about these classifications can be found in the Supplementary material S1.

#### Niki-TCGA dataset

The Niki-TCGA dataset is comprised of WSIs from two separate sources. 36 samples (25 NIFTP, 11 benign) were obtained from “Box A” of the [Nikiforov online repository](https://image.upmc.edu/NikiForov%20EFV%20Study/), which relates to research performed by Nikiforov et al. (31). The study accepted WSI contributions from 13 institutions across six different countries, before a panel of 24 expert thyroid pathologists determined each slide’s classification. The research sought to establish consensus diagnostic criteria for classifying NIFTP as a separate subcategory and therefore accepted many borderline cases which were considered difficult to diagnose even by expert pathologists (20, 31). Further information regarding the selection and designation of these samples can be found in Supplementary material S2. 31 PTC-like and 2 non-PTC-like samples (Table 2) were sourced from [The Cancer Genome Atlas (TCGA) Thyroid Carcinoma study](https://portal.gdc.cancer.gov/projects/TCGA-THCA). This study consists of 507 different patient samples, collected from 20 separate tissue source sites. Each patient was originally diagnosed with PTC, before a board-certified pathologist assigned fine-grained subtyping to each sample (43).

**Table 2.**
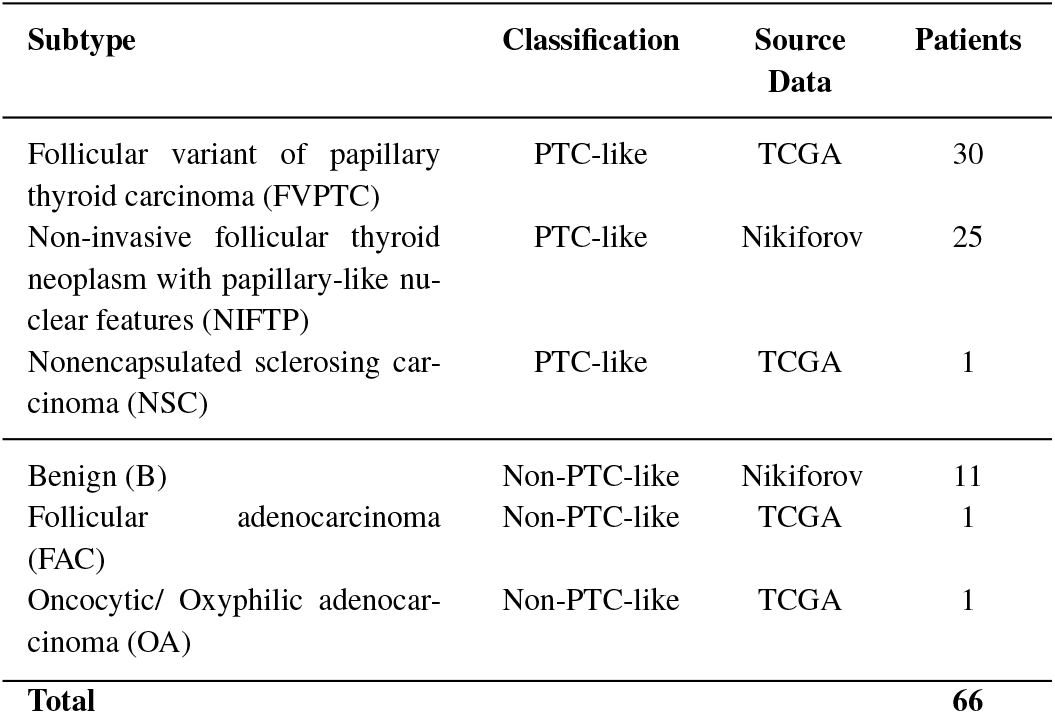
Sample subtypes present in the Niki-TCGA dataset, comprised of images sourced from the Nikiforov Box A samples as well as the TCGA Thyroid Carcinoma study. The table includes each sample’s classification as PTC-like or not and the number of patient samples in each category.

| Subtype | Classification | Source Data | Patients |
| --- | --- | --- | --- |
| Follicular variant of papillary thyroid carcinoma (FVPTC) | PTC-like | TCGA | 30 |
| Non-invasive follicular thyroid neoplasm with papillary-like nuclear features (NIFTP) | PTC-like | Nikiforov | 25 |
| Nonencapsulated sclerosing carcinoma (NSC) | PTC-like | TCGA | 1 |
| Benign (B) | Non-PTC-like | Nikiforov | 11 |
| Follicular adenocarcinoma (FAC) | Non-PTC-like | TCGA | 1 |
| Oncocytic/ Oxyphilic adenocarcinoma (OA) | Non-PTC-like | TCGA | 1 |
| **Total** |  |  | **66** |

The Niki-TCGA dataset was formed specifically to provide a robust test of a model’s generalizability. The FVPTC and NIFTP subtypes are commonly misdiagnosed by expert clinicians, and therefore are most important for an ML model to be able to predict. They also account for only 9 samples each within the T&T training dataset, mimicking the severe lack of training data which is often prohibitive to highly generalizable models in practice. Lastly, the dataset has high heterogeneity in terms of staining, resolution, and image quality, due to the combination of WSIs from multiple sources. Additionally, both data sources consist of patient samples gathered from diverse hospitals with varying demographics.

#### Niki-TCGA data pre-processing

WSIs for each sample in the dataset were downloaded, before a trained pathologist identified the neoplastic regions of interest within each slide which were indicative of the underlying classification (Supplementary material S3). From these regions, the OpenSlide function ‘DeepZoomGenerator’ was used to extract non-overlapping 512 × 512 px crops, before 20 from each sample were selected at random for inclusion within the dataset.

Figure 2 shows PTC-like and non-PTC-like examples from both the T&T and Niki-TCGA datasets, displaying the range of staining and resolution present.

**Fig. 2.**
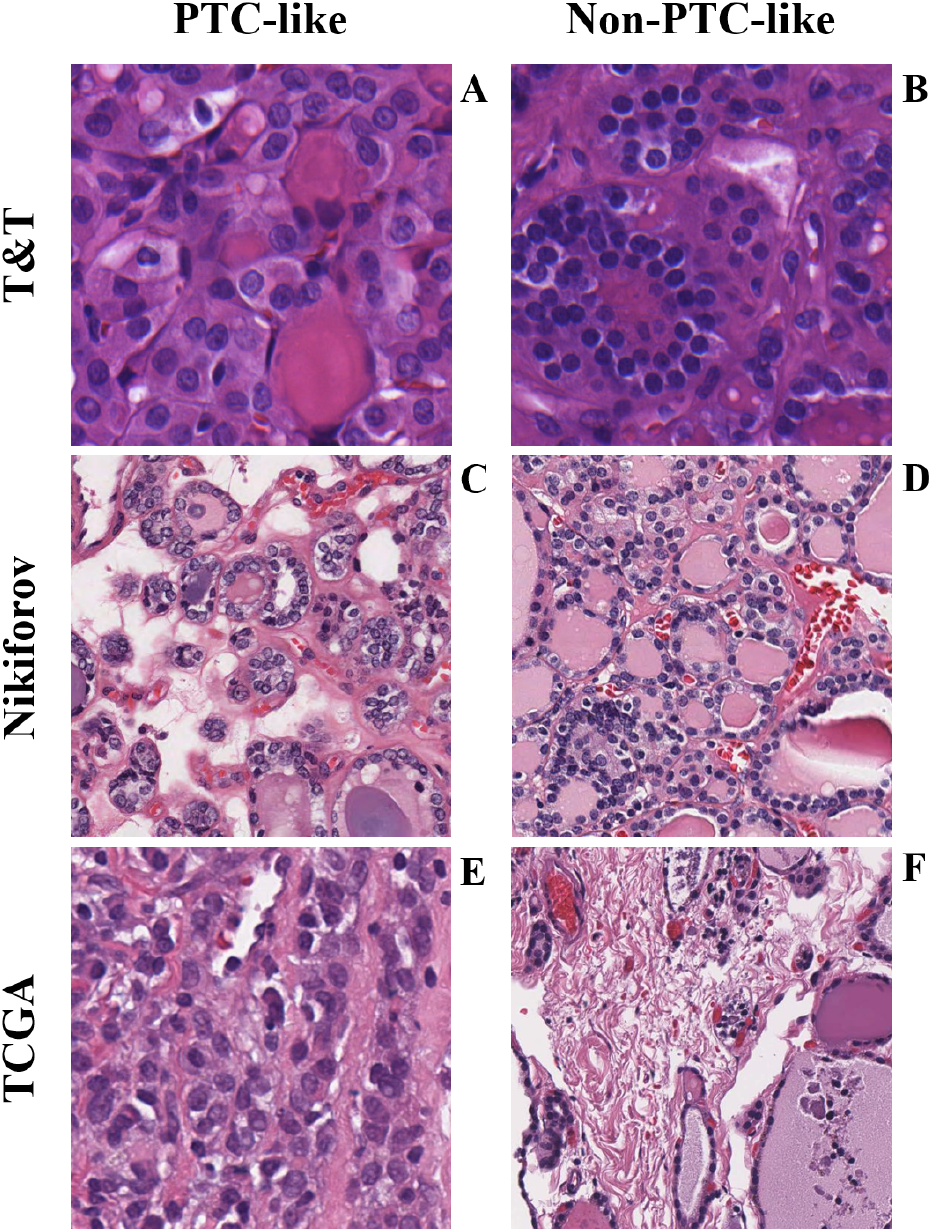
Example 512 × 512 px image crops taken from the three sources which comprise the T&T and the Niki-TCGA datasets. The classification and dataset identifier of each sample is as follows: (A) T&T - Papillary thyroid carcinoma (Dataset ID: 47h), (B) T&T - Follicular thyroid adenoma (Dataset ID: 39j), (C) Nikiforov - Noninvasive follicular thyroid neoplasm with papillary-like nuclear features (Dataset ID: A008), (D) Nikiforov - Benign (Dataset ID: A058), (E) TCGA - Papillary carcinoma, follicular variant (Dataset ID: TCGA-FY-A3I5), (F) TCGA - Oxyphilic adenocarcinoma (Dataset ID: TCGA-BJ-A192)

### Generative adversarial network (GAN)

GANs are comprised of two neural network models, referred to as the generator and the discriminator. The generator aims to learn to approximate the underlying distribution of the training data in order to produce high-quality synthetic images. The discriminator conversely learns to discern the difference between these GAN-generated fakes and the real images (32).

The training is adversarial by nature as the two networks compete to improve at their respective roles. An equilibrium is reached when the generator is producing synthetic images which are indistinguishable from the underlying real data, and thus the discriminator can make predictions with only 50% accuracy.

This two-player min-max game can be represented by the following value function:

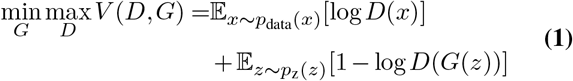

The StyleGAN2-ADA framework (44) was selected because it was specifically designed to produce high quality synthetic images from limited training data. It utilizes an extensive array of 18 transformations to augment the discriminator network’s training data. The method then adapts the level of augmentation based on feedback from an overfitting heuristic during model training.

The PyTorch implementation of StyleGAN2-ADA was adapted from the official StyleGAN2-ADA GitHub repository and trained with the following parameters:

- “cfg = paper512” to mirror the parameter settings used by Karras et al. (2020) (45) for the BRECAHAD dataset – a small dataset containing 162 breast cancer histopathology images (46).
- “cond = 1” ensures the GAN is trained conditionally using the labels provided, and so is subsequently able to produce images for a given class.
- “mirror = 1” includes x-flips of each image in the dataset, effectively doubling the training images.
- “kimg = 25000” trains the GAN for up to 25 million training steps. All GANs tested in the StyleGAN2-ADA paper were shown to produce their highest quality images before this point in training.

The GAN was trained with two different conditional labelling approaches to assess the quality and diversity of the synthetic images produced:

- Binary: training samples were labelled to be either PTC-like (1.0 label) or non-PTC-like (0.0).
- Multi-class: labelled according to their individual subtypes, being PTC (0.0 label), NIFTP (1.0), FVPTC (2.0), FA (3.0) and FTC (4.0). This enables the trained GAN to produce synthetic images of any given subtype.

#### GAN pre-processing and evaluation

The original images in the T&T dataset are 1916 × 1053 px. The GAN was trained to produce 512 × 512 px image patches, which are subsequently used to augment training data for a deep learning classifier (DLC) to classify images of the same size. To ensure that the maximum data was made available to train the GAN, the T&T images were split into equal 512 × 512 px patches with minimal overlap (Figure 3). This increased the number of training images from 1,496 (if only cropped centrally) to 12,038.

**Fig. 3.**
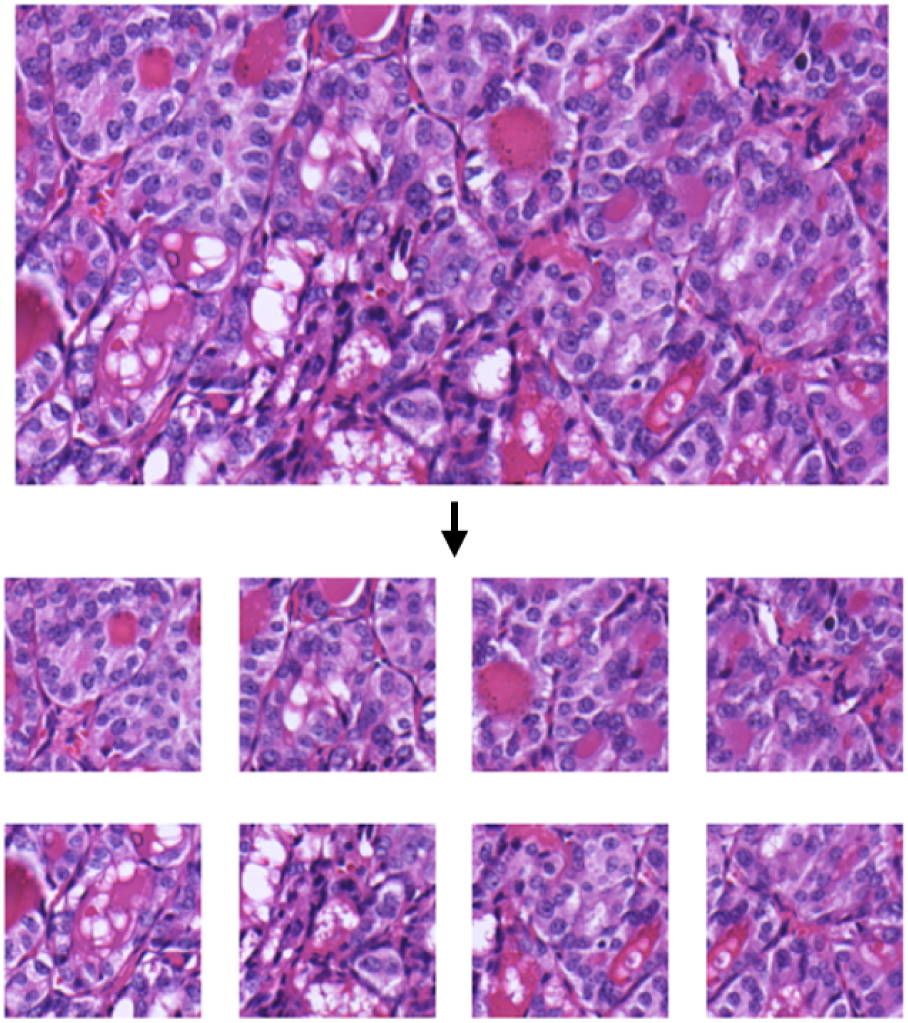
An example of one 1,916 × 1,053 px patient sample from the T&T dataset being split into 8 equal 512 × 512 px patches to be used for GAN training.

GAN training can be notoriously unstable, as the networks are prone to overfitting and mode collapse (40). To assess progression, batches of synthetic images were manually assessed at fixed intervals and compared to real images. Additionally, Fréchet Inception Distance (FID) (47) was used as the programmatic evaluation criteria for the Generator model. FID computes the Fréchet Distance (48) between multivariate Gaussian approximations of both the real and generated image distributions.

A lower FID score implies a greater alignment between the fake and real images and has been correlated with human judgement regarding the quality of the generated images (49). Despite this, the metric is not considered perfect, and several papers have warned against over-reliance on FID to assess GAN improvement (45, 49).

In this paper FID was used as the target metric for GAN training. Despite this, the true assessment of the quality of the generated images will be whether their addition to the real images during training improves the classification performance of the DLC on the unseen test data. This report therefore assesses the impact of GANs in an applied scenario, rather than purely evaluating the generative technique in isolation.

### Deep learning classification (DLC) model

The DLC model is based on the research performed by Bohland et al (2021) (20) using the T&T dataset. A pre-trained version of ResNet101 (50) was loaded using the Torchvision module, and the final output layer was altered to output a binary prediction – whether the image is PTC-like or not.

This model was then fine-tuned on the histopathology images in the T&T training set. The Albumentations package was used to apply random cropping, flipping, adaptive histogram equalization, blurring, Gaussian noise and Fourier Domain Adaptation (51) transformations.

A grid search found the following optimal parameters for the DLC:

- Initial learning rate of 1e-3.
- Learning rate decay of 5e-1 if the validation loss does not decrease for 10 epochs.
- Early stopping if the validation loss doesn’t decrease for 50 epochs.
- Optimizer weight decay of 1e-6.
- Maximum epochs of 500.

The Adam optimizer (52) and cross-entropy loss were also used. This setup utilized four GPUs and a batch size of 64 to obtain the most stable training results at a speed of 46 seconds per epoch.

Five-fold cross validation was used to evaluate the model’s performance on the T&T dataset. Each data split contained 60% for training, 20% for validation and 20% for testing. There are 156 patients in the T&T data, but each patient has multiple image slides associated with them. The accuracy metrics are therefore calculated at a patient level, rather than at a slide level. The final classification is determined by majority voting, in the case of a 50:50 split decision between the slides, the wrong class is assigned to the patient.

To evaluate the model’s generalizability to external data, the model is trained using 80% of the original T&T data, with the remaining 20% used as the validation set. The Niki-TCGA dataset is then used as the test set, with accuracy also calculated at the patient-level. F1 score is used as an additional metric due to the class imbalance in the external data. In both instances, if GAN-generated samples are used, they are included within the training data only.

### Synthetic data augmentation

Several different approaches to GAN-based augmentation were tested. The two most successful have been presented in this paper whilst full details can be found in Supplementary material S4.

“Binary 100” uses a GAN trained with binary classification labels (PTC-like or non-PTC-like) and increased the frequency of each of these binary class images by 100% in the training data. “MC 100” used a multi-class GAN, trained to produce images of every subtype in the T&T training data. This method augments the subtype with the most real images (FA) by 100%, and then equalizes all other subtypes to that level.

## Results

### GAN training

Table 3 shows the results of the StyleGAN2-ADA training. The FID score significantly improves to 5.05 when the original 1,916 × 1,053 px images were split into non-overlapping 512 × 512 px crops to increase the amount of available training data. This score is similar to the 4.67 average score that Karras et al. (2020) (45) obtained for three 5,000 image “Animal Faces (AFHQ)” datasets (Choi et al. 2020). It is also notably lower than the FID score of 15.71 Karras et al. achieved on the BreCaHAD dataset, which consisted of 162 binarily labelled breast cancer histopathology images, reorganized into 1,944 partially overlapping 512 × 512 px overlapping crops.

**Table 3.** Results from GAN training showing the minimum FID score under different approaches. Where data labels are “None” the GAN was trained to produce a generic thyroid histopathology image. “Binary” indicates the GAN has been trained conditionally on samples either labeled as PTC-like or Non-PTC-like. “Multi-class” denotes that the GAN was trained with conditional labels for each classification subtype - PTC, NIFTP, FVPTC, FA and FTC.

| Approach | Training Images | Data Labels | Min. FID |
| --- | --- | --- | --- |
| StyleGAN2-ADA | 1,496 | None | 15.39 |
| StyleGAN2-ADA | 1,496 | Binary | 18.38 |
| StyleGAN2-ADA | 12,038 | Binary | 5.10 |
| StyleGAN2-ADA | 12,038 | Multi-class | **5.05** |

Figure 4 illustrates synthetic images generated by GANs trained on binary labels. It compares images produced using two different training sets: one consisting of the 1,496 centrally cropped T&T images, producing an FID score of 18.38, and another consisting of 12,038 overlapping crops from the same T&T dataset, resulting in an FID score of 5.10.

**Fig. 4.**
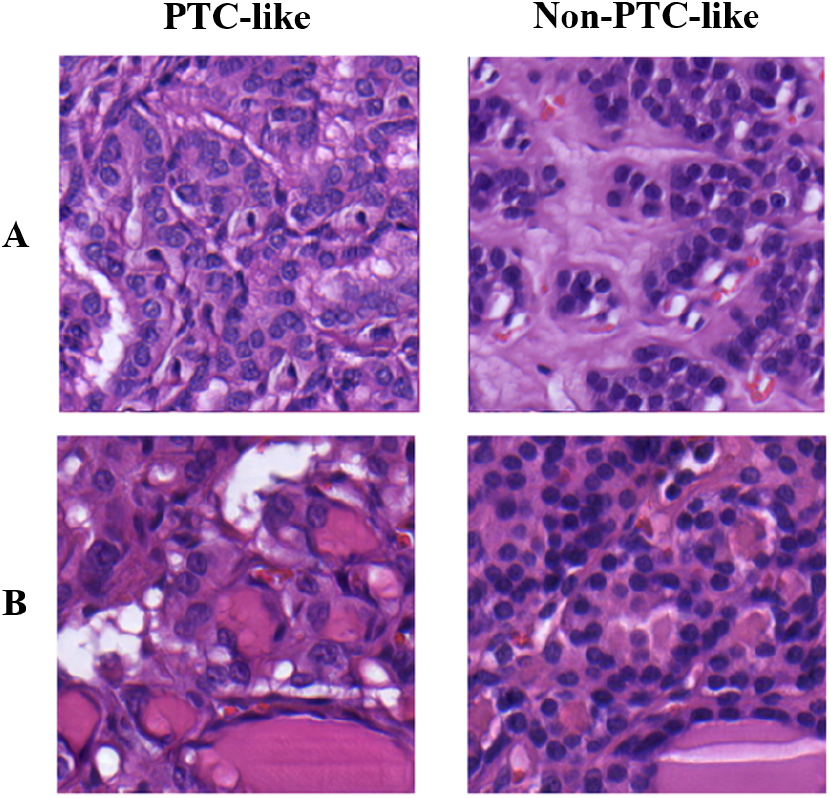
Examples of PTC-like and non-PTC-like images produced after Style-GAN2 ADA was trained conditionally using: (A) the original 1,496 T&T dataset with 512 × 512 px crops extracted from the centre of each image, achieving an FID score of 18.38. (B) Compared with the expanded T&T dataset which extracted 12,038 overlapping 512 × 512 px crops from the original images, resulting in an FID score of 5.10.

Figure 5 presents a comparison between synthetic images from each subtype designation generated by the multi-class GAN and real images from the T&T data.

**Fig. 5.**
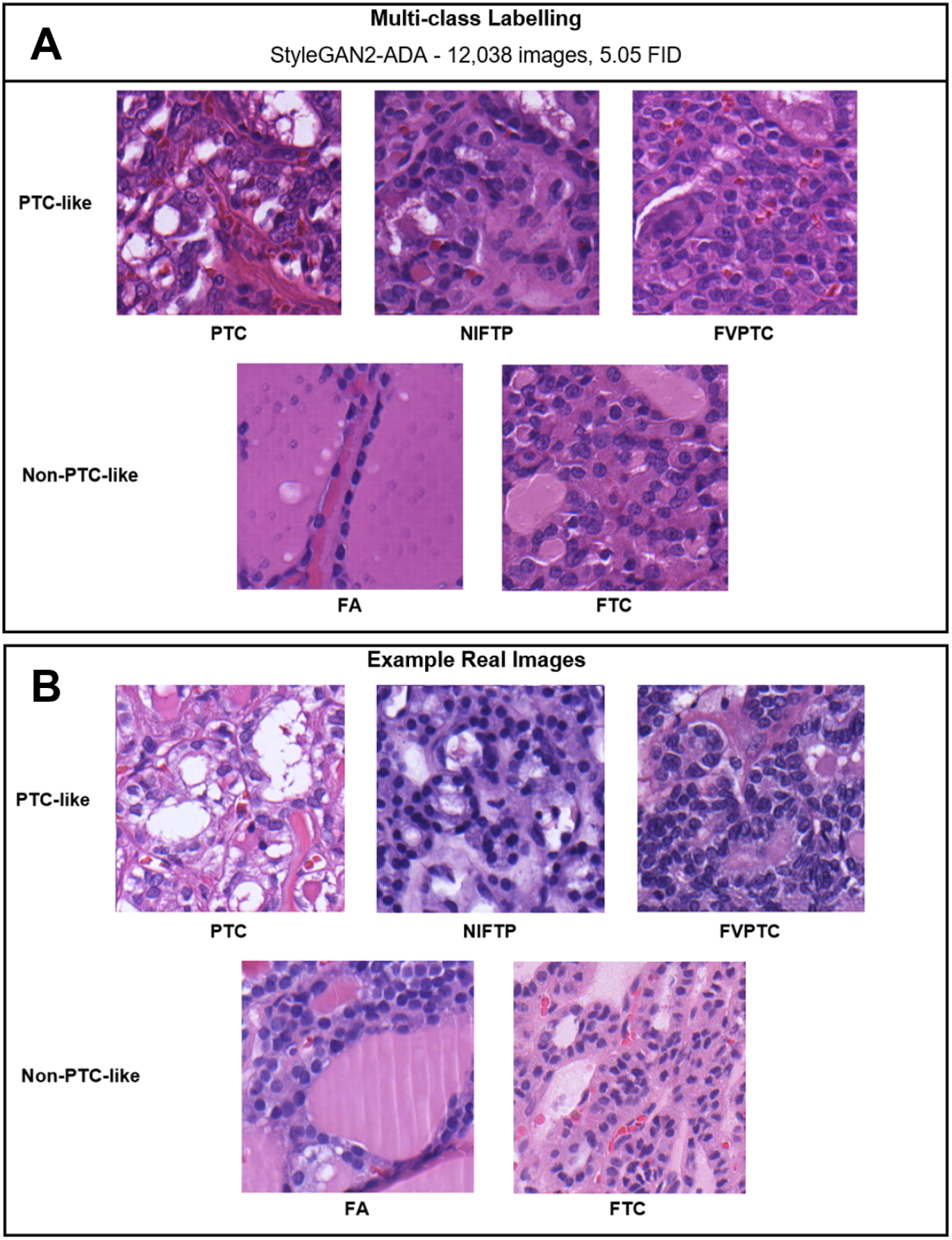
(A) Example 512 × 512 px images for each class subtype produced by the multi-class conditional GAN. (B) Comparison real image crops taken from the T&T dataset for each class subtype.

### DLC results

#### T&T dataset

Table 4 displays the five-fold cross-validation performance of the DLC model trained to classify images from the T&T dataset. The baseline model achieved an average overall accuracy of 83.97%. The model performs notably better on the majority classes (PTC, FA and FTC), with an average accuracy of 88.7%. However, the minority subtypes (NIFTP and FVPTC), which are considered more borderline diagnoses, are classified correctly with 49.95% average accuracy.

**Table 4.** Five-fold cross-validation fold accuracy metrics for DLC models trained to predict samples as PTC-like or Non-PTC-like. The training of each fold is performed with a 60% split of the data used for training, 20% for validation and 20% for testing. Baseline includes no GAN-generated images within the training data, whilst Binary 100 and MC 100 include synthetic data augmentation as set out in Synthetic data augmentation.

| Approach | PTC-like |  |  | Non-PTC-like |  | Overall Acc. |
| --- | --- | --- | --- | --- | --- | --- |
|  | PTC | FVPTC | NIFTP | FA | FTC |  |
| **Baseline** | 83.02% | 55.56% | 44.44% | 92.45% | **90.63%** | 83.97% |
|  | (44/53) | (5/9) | (4/9) | (49/53) | (29/32) | (131/156) |
| *Std. Dev.* | 13.37% | 35.35% | 41.83% | 7.67% | 13.08% | 5.85% |
| **Binary 100** | **90.57%** | **77.78%** | 55.56% | **98.11%** | 81.25% | 88.46% |
|  | (48/53) | (7/9) | (5/9) | (52/53) | (26/32) | (138/156) |
| *Std. Dev.* | 5.76% | 40.00% | 44.72% | 4.00% | 13.85% | 6.50% |
| **MC 100** | **90.57%** | **77.78%** | **88.89%** | 96.23% | 84.36% | **90.38%** |
|  | (48/53) | (7/9) | (8/9) | (51/53) | (27/32) | (141/156) |
| *Std. Dev.* | 9.96% | 40.00% | 20.00% | 4.45% | 15.68% | 4.56% |

The Binary 100 GAN augmentation strategy increased the overall classification accuracy by 5.3%, from 83.97% to 88.46%. The approach particularly benefitted the PTC and FA classes, which increased in accuracy by 9.1% and 6.1% respectively. Despite this, FTC samples were classified with 10.3% less accuracy.

The MC100 approach resulted in the highest accuracy classifier (90.38%), 7.6% greater than the baseline approach. Overall, the MC100 model predicts the minority class subtypes (FVPTC and NIFTP) with 83.33% accuracy, a notable improvement over the baseline of 49.95%. It is also the method with the lowest standard deviation across cross-validation folds, implying it is the most consistent classifier.

#### Niki-TCGA dataset

Similar to the five-fold cross-validation results seen with the T&T data, augmenting the training data with GAN-generated samples has a positive impact on the performance of the DLC when tested on the external Niki-TCGA data (Table 5). Again, MC100 was the best performing approach, increasing overall patient accuracy by 35.71% and F1 score by 42.30% over the baseline classifier.

**Table 5.** DLC results when tested on the Niki-TCGA dataset. The “Baseline” model utilizes 80% of the T&T dataset for training and 20% for validation. The “Binary 100” and “MC 100” models both use GAN-generated synthetic samples during training (described in Synthetic data augmentation), to augment the original T&T training data.

| Approach | PTC-like |  |  | Non-PTC-like |  |  | Accuracy | F1 Score |
| --- | --- | --- | --- | --- | --- | --- | --- | --- |
|  | FVPTC | NIFTP | NSC | B | FAC | OA |  |  |
| **Baseline** | 46.67% | 36.00% | **100.00%** | 36.36% | 0.00% | 0.00% | 40.58% | 35.13 |
|  | (14/30) | (9/25) | (1/1) | (4/11) | (0/1) | (0/1) | (28/69) |  |
| **Binary 100** | 60.00% | 40.00% | 0.00% | 54.55% | **100.00%** | 0% | 50.72% | 45.69 |
|  | (18/30) | (10/25) | (0/1) | (6/11) | (1/1) | (0/1) | (35/69) |  |
| **MC 100** | **63.33%** | **44.00%** | 0.00% | **72.73%** | 0.00% | 0.00% | **55.07%** | **49.99** |
|  | (19/30) | (11/25) | (0/1) | (8/11) | (0/1) | (0/1) | (38/69) |  |

## Discussion

In this work we successfully trained a GAN which could produce high quality synthetic thyroid histopathology images using a limited dataset of images from 156 patients. The GAN FID scores were comparable to prior state-of-the-art results for similar-sized datasets from different domains (45), showing the applicability of the StyleGAN2-ADA approach to the medical imaging domain.

We demonstrated that including these synthetic samples in the training data for a DLC resulted in a more robust model which could generalize more effectively to external data. This experimental setup mirrored the key challenges present when considering deploying in-silico models in practice, focusing on the real-world issues of both data scarcity and heterogeneity. Our results support the use of GANs as a partial solution, offering evidence that generative models can learn some of the key characteristics of biological image data by approximating the underlying population distribution, improving classification performance in the absence of more real training images.

Several GAN-augmentation strategies were trialled on two test sets – T&T (via five-fold cross-validation) and the Niki-TCGA external data. In both instances, augmenting the training data with GAN-generated synthetic samples proved to be beneficial. The narrowing of the domain gap between the image datasets, as a result of including synthetic samples, can be observed in Supplementary material S5.

The PTC samples in the T&T test data were classified with an accuracy of 90.57%, an increase of 9.1% over the baseline model, which uses no synthetic images during training. PTC is the most common thyroid malignancy and has the most clear and numerous distinguishing nuclear features. Representing these features would therefore be most straightforward for the GAN to learn, resulting in synthetic images which clearly improved the DLC training process.

The minority PTC-like subtypes, FVPTC and NIFTP, are much more difficult for a pathologist, or an ML model, to identify because the PTC-like features can be found within encapsulated follicular lesions, are less numerous and distinct, and are often surrounded by benign-appearing cells.

For these subtypes, using a multi-class GAN was most beneficial. In the case of the T&T data, the multi-class GAN achieved an average accuracy of 83.33%, compared to the binary GAN augmentation approach which yielded 66.68% accuracy, and the baseline accuracy of 49.95%. It is particularly impressive that the GAN was able to represent these subtype distributions effectively, given it was trained on only nine patient samples per minority class.

For non-PTC-like T&T samples (FA and FTC), the improvement from GAN-augmentation was not as clear-cut. For FA samples, which definitively lack any PTC-like nuclear features, the binary and multi-class GAN both improved classification accuracy by 6.1% and 4.1% respectively. This aptitude for correctly classifying FA class samples was again evident when testing on the Niki-TCGA dataset, as the GAN augmentation notably improved the DLCs ability to predict the benign samples, also definitively lacking the PTC-like features.

However, for the FTC class, classification accuracy decreased with both strategies, from 90.63% accuracy seen with the baseline model, to 81.25% for a binary GAN and 84.38% for a multi-class GAN. FTC samples are defined by a lack of nuclear atypia required for a PTC-like diagnosis. There is significant disagreement amongst pathologists about what level of nuclear atypia constitutes this diagnosis boundary and this can lead to significant inter-observer variability.

For the GAN conditioned on binary labels, the non-PTC-like images it produced likely aligned more closely with the FA distribution, which is the majority non-PTC-like class. This resulted in higher accuracy when testing on FA samples (98.11%), but a relatively worse performance on FTC samples (81.25%).

In the case of the multi-class GAN augmentation, the number of PTC-like samples with less distinct PTC-like features (i.e., the FVPTC and NIFTP classes) in the training data are substantially increased. This likely made the DLC better at identifying these features compared to the baseline model, resulting in lower sensitivity of the model for the FTC class specifically.

Results for the external Niki-TCGA dataset provided evidence that the DLC was not overfitting on these subtypes and was able to generalize with increased accuracy. Again, MC100 was the best performing augmentation strategy, improving both classification accuracy and F1 score compared with the baseline model. The consistency of results across test sets provides additional confidence in the reliability of the approach and its potential applicability to similar scenarios in different medical imaging contexts.

Whilst the combined accuracy of 55% (30/55) for the NIFTP and FVPTC samples in the Niki-TCGA data is not high, it is 31.6% greater than the 41.8% (23/55) accuracy achieved by the baseline. This study did not aim to identify the optimal classifier for this data but, instead, attempted to isolate the impact of GAN-generated synthetic training samples on a model’s ability to adapt to histopathological data from different domains. For future work it would be interesting to investigate whether more advanced deep learning-based architectures, such as the Swin Transformer (53) or ConvNeXt (54), would benefit to a greater or lesser extent from the inclusion of these synthetic examples.

An alternative solution to bridging the domain gap between datasets could be to use a style-transfer GAN. This approach aims to separate images into “content” and “style” spaces (55). In the context of this report, content would represent the various structural features of the histopathology images, whilst the domain differences, such as the stain colour or resolution, would be considered the style.

Finally, one criticism of deep learning architectures is that they function like “black box” models. Explainable AI is particularly important in healthcare as accountability, trust and understanding bias are integral components to the functioning of the system as a whole (56). Using a self-attention GAN (SAGAN) (57) could partially alleviate these issues. The attention mechanism (58) can be visualized to provide feedback regarding which parts of the image were most important for the model’s classification decision. This could form part of an important feedback loop for a pathologist to understand and interpret the model output, increasing trust in the its ability to accurately classify pathological or other medical images.

## Acknowledgements

The authors would like to thank Ryan Reavette for his guidance and feedback during the project, as well as the initial inspiration for the topic on GANs. We would also like to thank Jamie Holdstock for his contribution towards developing ‘ripsvs’, a tool to download image patches from online-hosted Aperio ImageScope WSIs.

## Data and source code availability

The data underlying the findings of this study, where not already publicly accessible, are available on request from the corresponding author. Source codes, as well as underlying data, are freely available at: https://github.com/williamdee1/ThyCa-GAN.

## Funding

The authors received no financial support for the research, authorship, and/or publication of this paper.

## Conflicts of Interest

None declared.

## Supplementary Information

### S1. Additional information regarding subtype classifications

#### Follicular cell-derived neoplasms

Benign thyroid tumours can be classified into several types based on their histopathological features. The most common types of benign thyroid tumours include:

- **Follicular adenoma (FA):** FA is derived from follicular cells of the thyroid gland. It is characterized by the presence of well-defined follicles that are lined by a single layer of cuboidal epithelial cells. These tumours are usually encapsulated and can be difficult to distinguish from normal thyroid tissue (59).
- **Follicular adenoma with papillary architecture:** This tumour is usually encapsulated with intra-follicular centripetal papillary growth, and the lesional cells lack nuclear features of papillary thyroid carcinoma (PTC) (60).
- **Oncocytic adenoma of the thyroid:** Both benign and malignant oncocytic thyroid tumours exhibit papillary architecture and are composed of oncocytes, which are large, eosinophilic cells with granular eosinophilic cytoplasm (61).

Overall, benign thyroid tumours are usually well-circumscribed and show a relatively uniform population of cells with no evidence of invasion or metastasis. However, it is important to note that some benign tumours can have atypical features that may require careful monitoring or even surgical removal. It is worth noting that although these tumours are generally considered benign, there is always a risk that they could progress to malignancy.

#### Malignant thyroid tumours

- **Papillary thyroid carcinoma (PTC):** PTC is the most common type of thyroid cancer, accounting for approximately 80% of cases. It arises from the follicular cells of the thyroid gland (62). The histopathology of PTC typically shows well-differentiated papillary structures with a fibrovascular core (hallmark of PTC). These structures are lined by one or several layers of neoplastic epithelial cells with crowded nuclei. The nuclei may display irregular contours, nuclear pseudo-inclusions and nuclear membrane irregularities (nuclear grooves). The chromatin is characterised by chromatin clearing, margination and glassy nuclei (31, 60, 63). In addition, PTC may exhibit psammoma bodies, which are concentrically laminated calcific nodules that represent the ghosts of dead papillae. These are a characteristic finding in PTC (63). In advanced cases, the cancerous cells can invade through the thyroid capsule, blood vessels, and lymphatics. Overall, the histopathology of PTC is characterized by well-differentiated papillary structures with neoplastic epithelial cells displaying nuclear changes that are considered a diagnostic hallmark of this tumour entity and this is shown in the World Health Organization (WHO) classification (64).
- **Papillary thyroid carcinoma-like tumours (PTC-like):** PTC-like tumours are a group of neoplasms that are histologically similar to PTC but are not true PTCs. They are considered a type of follicular cell-derived thyroid neoplasm. These tumours are typically characterized by the presence of papillary projections and nuclear features that are closely resembling PTC. However, unlike PTC, these tumours do not have the characteristic growth pattern and are not confined to the thyroid gland. Some examples of PTC-like tumours include:
  – **Non-invasive follicular thyroid neoplasm with papillary-like nuclear features (NIFTP):** NIFTP is a noninvasive tumour that was previously classified as a variant of PTC. However, it is now recognized as a distinct entity by the WHO (64). NIFTP is diagnosed based on specific histopathological features which include the presence of follicular architecture, absence of invasion, and nuclear features resembling those seen in PTC but less prominent (65). NIFTP is considered to have a very low risk of recurrence or progression to malignancy and is therefore classified as a low-risk neoplasm (64). It is important to distinguish NIFTP from PTC which have a higher risk of recurrence and require more aggressive treatment.
  – **Follicular variant of papillary thyroid carcinoma (FV-PTC):** FV-PTC is a subtype of PTC. The histopathology of FV-PTC is characterized by the presence of follicular architecture and nuclear features typical to PTC. In addition to the typical nuclear features, FV-PTC may also show areas of vascular invasion and perineural invasion, which are associated with a higher risk of recurrence and metastasis (60, 64). The tumour is usually well-circumscribed and encapsulated (60). A definitive diagnosis of FV-PTC requires the presence of nuclear features of PTC in the follicular architecture.
  – **Nonencapsulated sclerosing carcinoma or diffuse sclerosing variant of papillary thyroid carcinoma (DSV-PTC):** A rare subtype of PTC, accounting for less than 1% of all thyroid malignancies. It is characterized by diffuse infiltration of the thyroid gland by tumour cells, resulting in a diffusely enlarged gland with no discrete nodules. Histologically, DSV-PTC is characterized by the presence of a sclerotic stroma, which is composed of fibrous tissue and lymphocytes. The tumour cells are arranged in a diffuse and infiltrative pattern, with no well-defined follicles or papillae. The nuclei of the tumour cells exhibit nuclear features of PTC. Psammoma bodies may be present within the tumour (66). DSV-PTC is often associated with lymph node metastases and a higher incidence of extrathyroidal extension compared to other types of PTC. It has also been reported to have a more aggressive clinical course and a higher recurrence rate, although this is still a matter of debate (66).
  In summary, PTC-like tumours are a group of neoplasms that are histologically similar to PTC but have different growth patterns and clinical characteristics. The treatment and prognosis of these tumours may vary depending on the specific subtype.
- **Follicular thyroid carcinoma (FTC)** is the second most common type of thyroid cancer, accounting for approximately 10-15% of all cases. FTC is derived from the follicular cells of the thyroid gland (67, 68). FTC shows a characteristic pattern of follicular growth, lined with neoplastic cells without the nuclear features of PTC (68). The histopathology of FTC is indistinguishable from follicular adenoma (FA) but the former appears more cellular and with irregular thick capsule (59). Histologically, FTC is distinguished from FA by the presence of capsular or vascular invasion. The presence of invasion is usually confirmed by examining the tumour capsule or the surrounding thyroid tissue for the presence of tumour cells. If invasion is identified, the diagnosis of FTC is made. In addition, FTC may show vascular invasion, with tumour cells present within blood vessels. This is another important diagnostic criterion for malignancy (59). Overall, the histopathology of FTC is characterized by the presence of follicular growth patterns, uniform neoplastic follicular cells, and invasion of the fibrous capsule and/or blood vessels being the hallmark feature that distinguishes it from FA.
- **Oncocytic carcinoma of the thyroid:** The tumour cells are identical to the benign counterpart. The tumour cells may exhibit the nuclear features of PTC or may have pleomorphic and hyperchromatic nuclei. Oncocytic carcinoma is associated with aggressive clinical behaviour in comparison to PTC (61, 69).

#### WHO Classification (2022) - Follicular cell–derived neoplasms

Supplementary material, Figure 1 contains a table of the WHO (2022) designation of the following PTC-like and Non-PTC-like subclasses:

- **PTC-like:** NIFTP, PTC, FV-PTC, DSV-PTC
- **Non-PTC-like:** Benign, FTC, Oncocytic/Oxyphilic carcinoma of the thyroid

**Supplementary Material, Figure 1.**
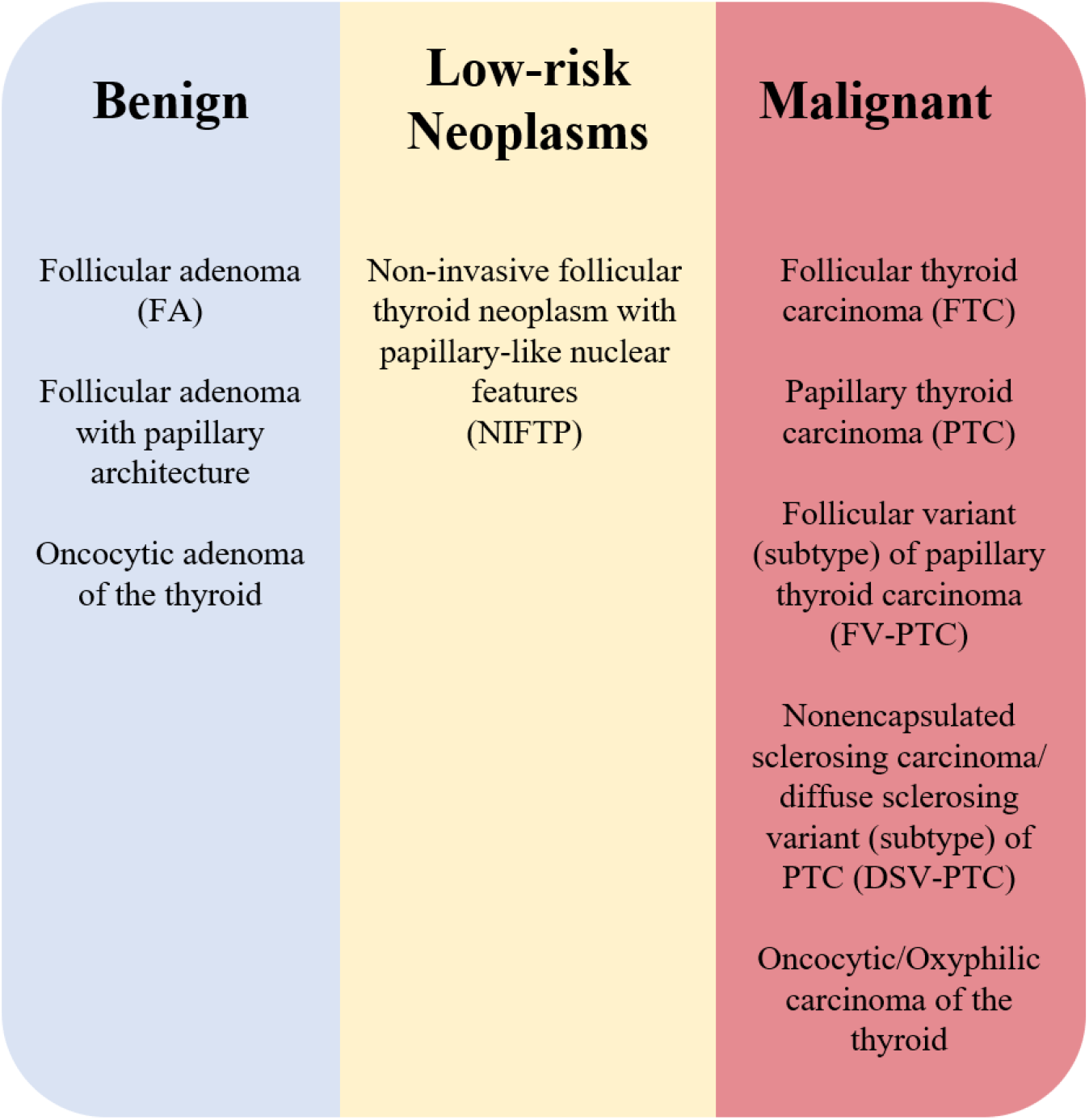
WHO (2022) designation of subclasses which appear in this report.

### S2. Niki-TCGA dataset selection

The “Nikiforov” dataset used by (20) consisted of 138 patient samples (103 PTC-like, 30 non-PTC-like) taken from “BoxA” of the Nikiforov online repository. The labels of these samples are neither shared by the reference paper nor are in the repository. Therefore, the designation of PTC-like or non-PTC-like could only be determined for 55 samples which were explicitly scored within Tables 5 and 6 of the Nikiforov supplementary materials (31). Of these, 36 samples remain uploaded in BoxA. 25 are designated as NIFTP (PTC-like) and 11 are benign (Non-PTC-like). These were included to form part of the external dataset used by this paper for testing model generalizability.

Additional thyroid histopathology samples were sourced from The Cancer Genome Atlas (TCGA) Thyroid Carcinoma study. The study contains the following samples:

- Papillary adenocarcinoma (PA) – 356 patients
- Follicular variant of papillary thyroid carcinoma (FVPTC) – 105 patients
- Papillary carcinoma, columnar cell (PCC) – 38 patients
- Nonencapsulated sclerosing carcinoma (NSC) – 4 patients
- One patient sample of: follicular thyroid carcinoma (FTC), follicular adenocarcinoma (FAC), and oxyphilic adenocarcinoma (OA)

From this, 30 FVPTC samples were selected as PTC-like samples for the Niki-TCGA dataset. This ensured a similar number of both minority classes (NIFTP and FVPTC) in the T&T training data would be present in the external test data. All non-PTC-like patient samples from TCGA were downloaded, however only three files were uncorrupted and could be included for testing.

The full Niki-TCGA dataset is summarized in Table 2. The dataset contains no samples belonging to the majority classes used to train the deep learning model (PTC, FA, and FTC), as well as subtypes that the model has never seen before – B, FAC, NSC and OA samples.

### S3. Neoplastic region identification

Supplementary material, Figure 2 shows two whole slide images after they have been annotated. From these identified regions of interest, 20 random image patches were extracted for each sample and used to form the Niki-TCGA dataset.

**Supplementary Material, Figure 2.**
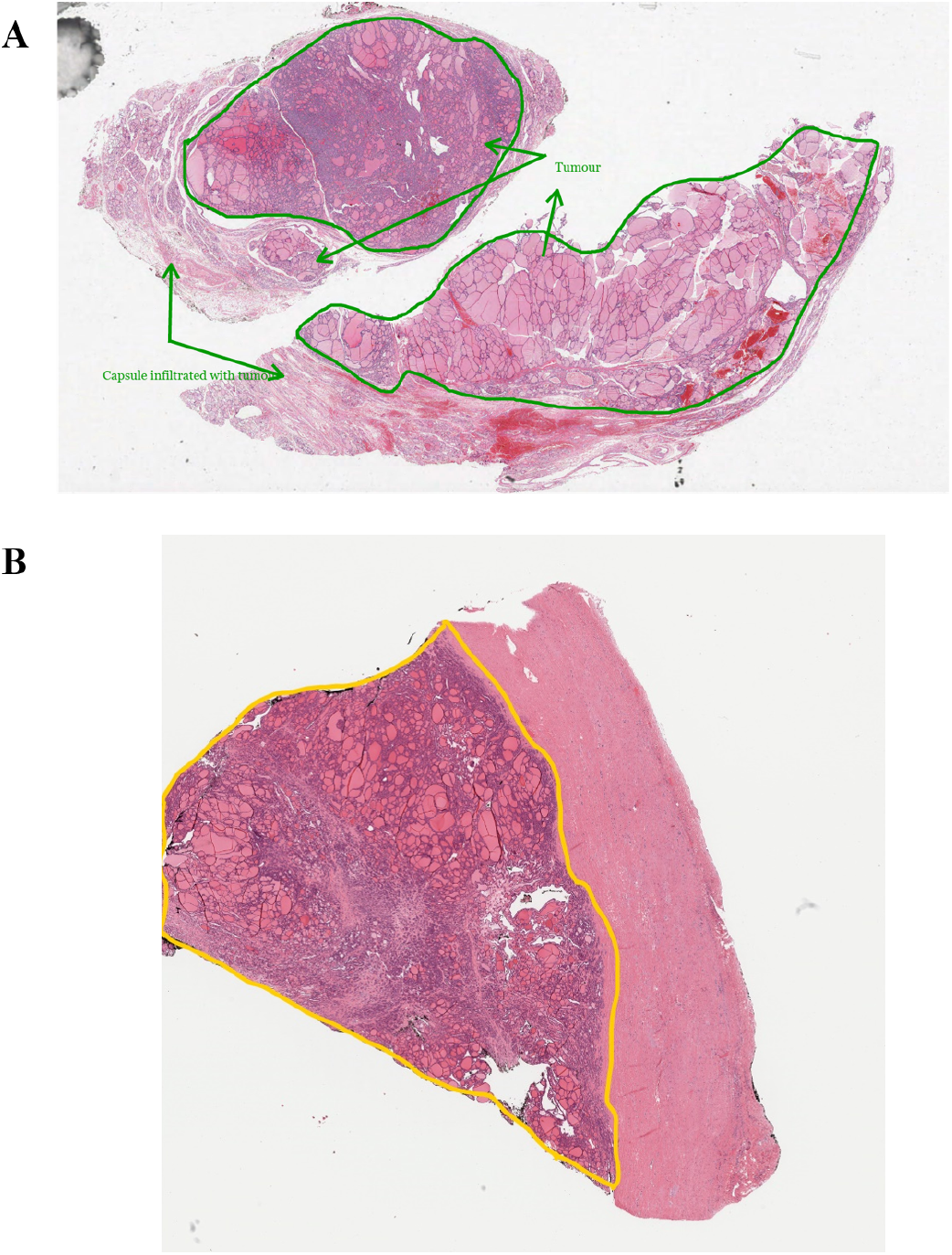
Whole slide images where neoplastic regions of interest corresponding to the classification designation have been identified. (A) corresponds to the NIK-A036 NIFTP sample from Box A of the online Nikiforov repository. (B) relates to the TCGA Thyroid Carcinoma study FVPTC sample TCGA-EM-A4G1.

### S4. GAN augmentations

This section includes details of the various GAN augmentation strategies. Supplementary material, Figure 3 and Figure 4 show the number of GAN synthetic images added to each class according to the strategy selected, the type of GAN trained and the data the model will be tested on.

**Supplementary Material, Figure 3.**
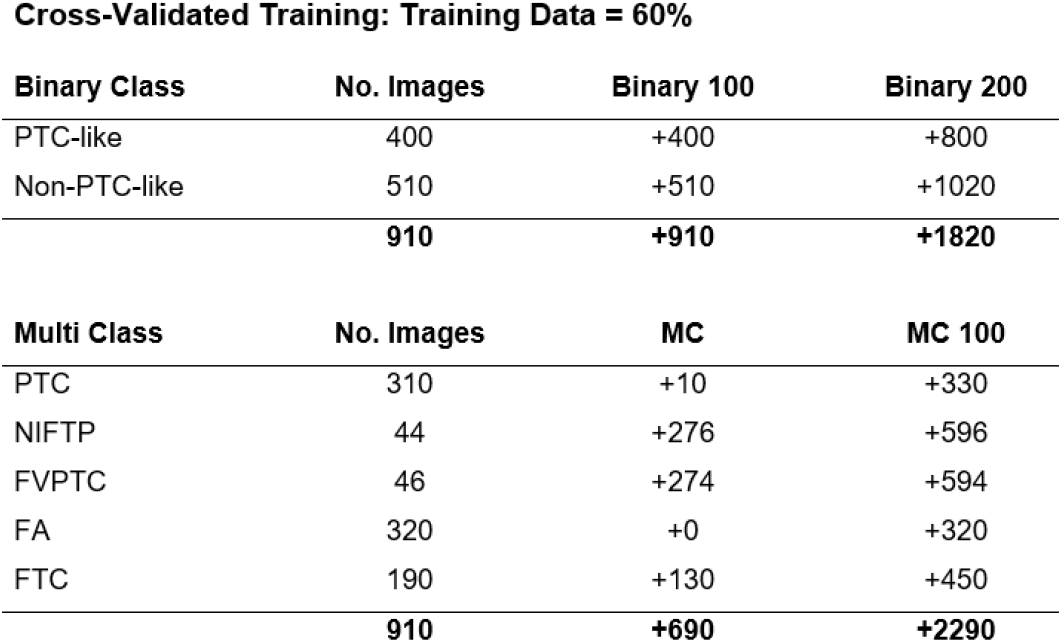
Table showing how many GAN-generated images are added to each classification type for each GAN augmentation strategy. In the case of cross-validated training, only 60% of the data is used for training the model, so the GAN augmentations are based off those 910 original real images. When using a binary-conditioned GAN the samples added are either designated to be PTC-like or Non-PTC-like. If the GAN was trained conditionally, then samples are added according to individual subtypes.

**Supplementary Material, Figure 4.**
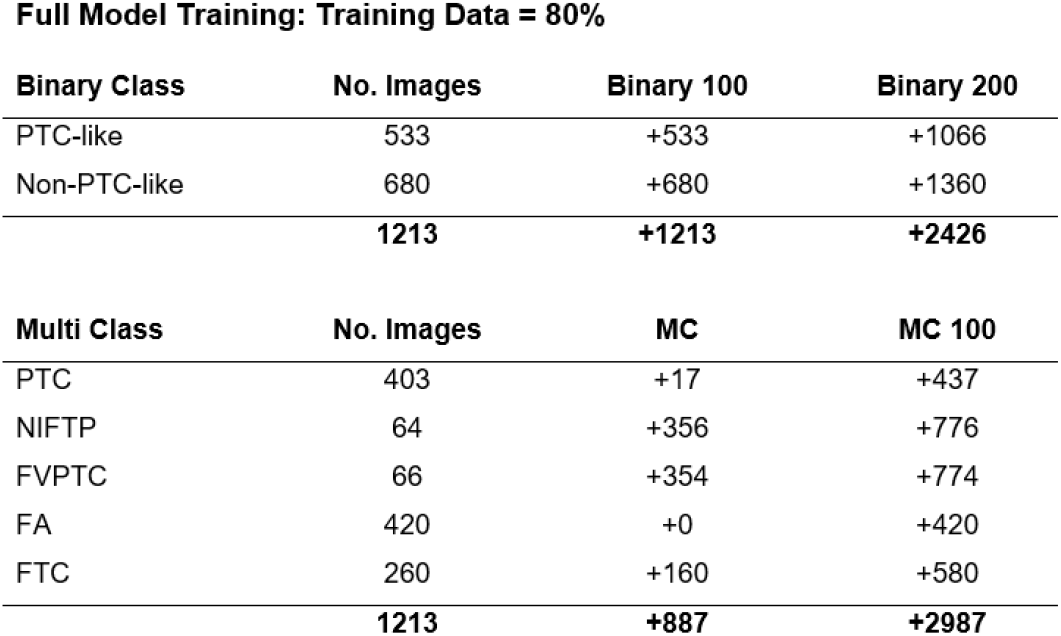
Table showing how many GAN-generated images are added to each classification type for each GAN augmentation strategy. When the model will be tested on the Niki-TCGA data, 80% of the T&T data is used to train the model, therefore the GAN augmentations are based off those 1,213 original real images. When using a binary-conditioned GAN the samples added are either designated to be PTC-like or Non-PTC-like. If the GAN was trained conditionally, then samples are added according to individual subtypes

### S5. Domain gap visualization

Supplementary material, Figure 5 (A) shows the T&T dataset images (the “Source Data”) as well as both the Nikiforov and TCGA images after they have been passed through a ResNet50 classifier and reduced to a two-dimensional representation by UMAP. Like the work of (20), there is a noticeable separation between the distributions of the datasets, with the Nikiforov and TCGA data being more closely aligned with each other than the T&T data, which is clearly distinct. Visualizing the disparity between the data distributions provides an insight into how difficult it will be for a model trained on one dataset to generalize to another, implying that a model trained on the T&T source data alone would generalize poorly to the Niki-TCGA data.

Supplementary material, Figure 5 (B) shows how this domain gap is influenced by the inclusion of 1,500 (750 PTC-like, 750 Non-PTC-like) GAN-generated synthetic images. The synthetic images largely overlap with the source T&T data; however, their addition has brought all the distributions closer together, thereby narrowing the domain gap between the datasets. This appears to be especially true for the TCGA images (green), which now lie much closer to the training data (which includes both the T&T data as well as the GAN-generated images).

Based on these visualizations, it appears likely that including the synthetic samples during model training will have a positive impact on the ability of the model to generalize to the Niki-TCGA data. It also implies that the TCGA samples would be predicted with greater accuracy compared to the Nikiforov samples, based on the proximity of the distributions. This was confirmed by our results in Table 5.

**Supplementary Material, Figure 5.**
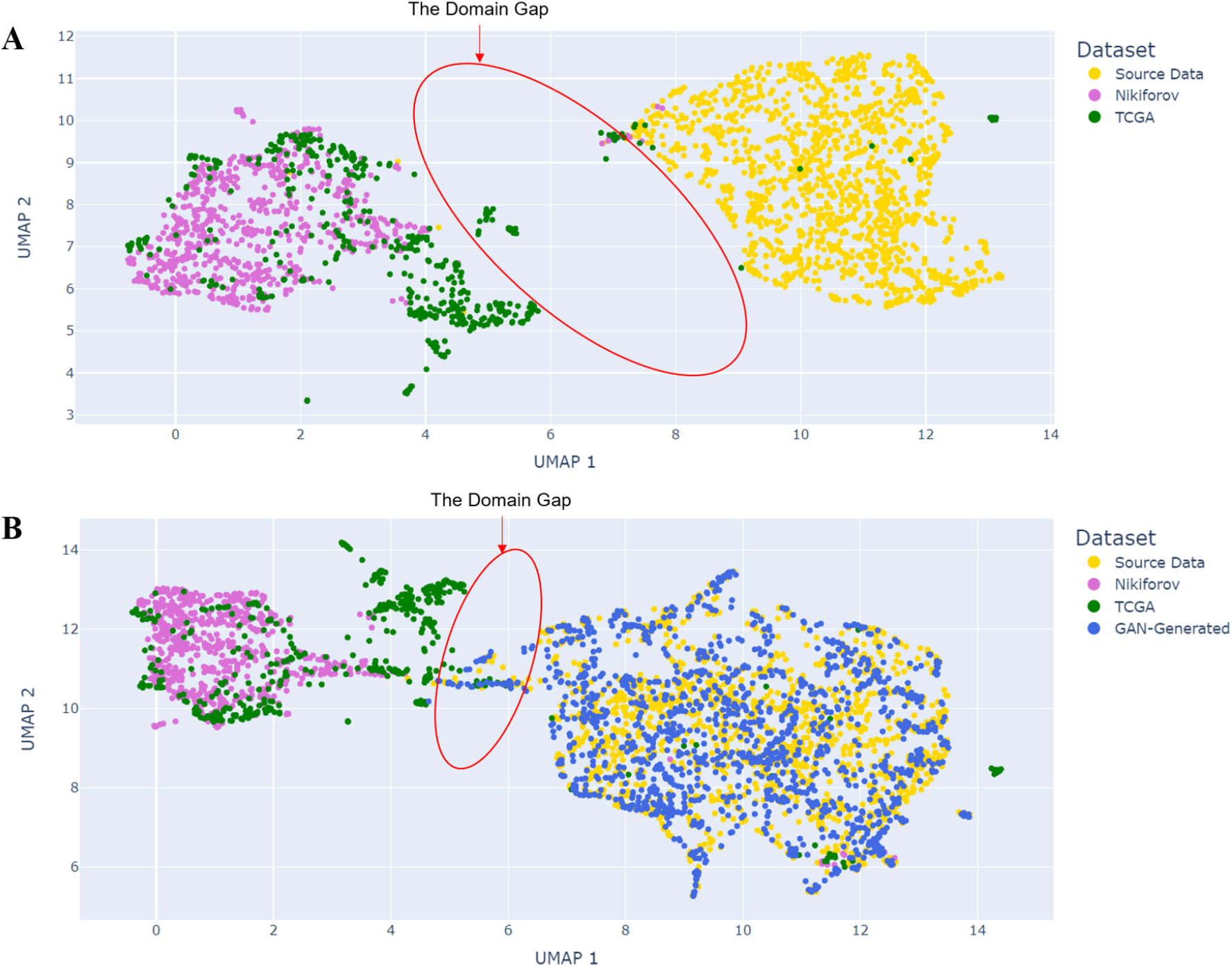
(A) A visualization of the domain gap which exists between the T&T (“Source”) dataset and that of the external test data, comprised of both TCGA and Nikiforov samples. The images were plotted after being fed through a ResNet50 classifier before being reduced to a two-dimensional representation by UMAP. (B) A plot of the UMAP two-dimensional representation of the T&T (“Source”), Nikiforov, TCGA and 1,500 GAN-generated synthetic samples. The domain gap has been annotated to reflect the distance in this two-dimensional space that largely separates the training data (T&T plus GAN images) from the test data (Nikiforov and TCGA).

### S6. Five-fold cross-validation results

When performing cross-validation there are *n* splits of the data according to how to many cross-validation folds there are. Included within this section are the results for each split of the five cross-validation folds tested on the T&T dataset. The overall results of each approach have been included within the main text, but the variation amongst subtypes and between folds is included here for completeness given the small dataset size.

**Supplementary Material, Figure 6.**
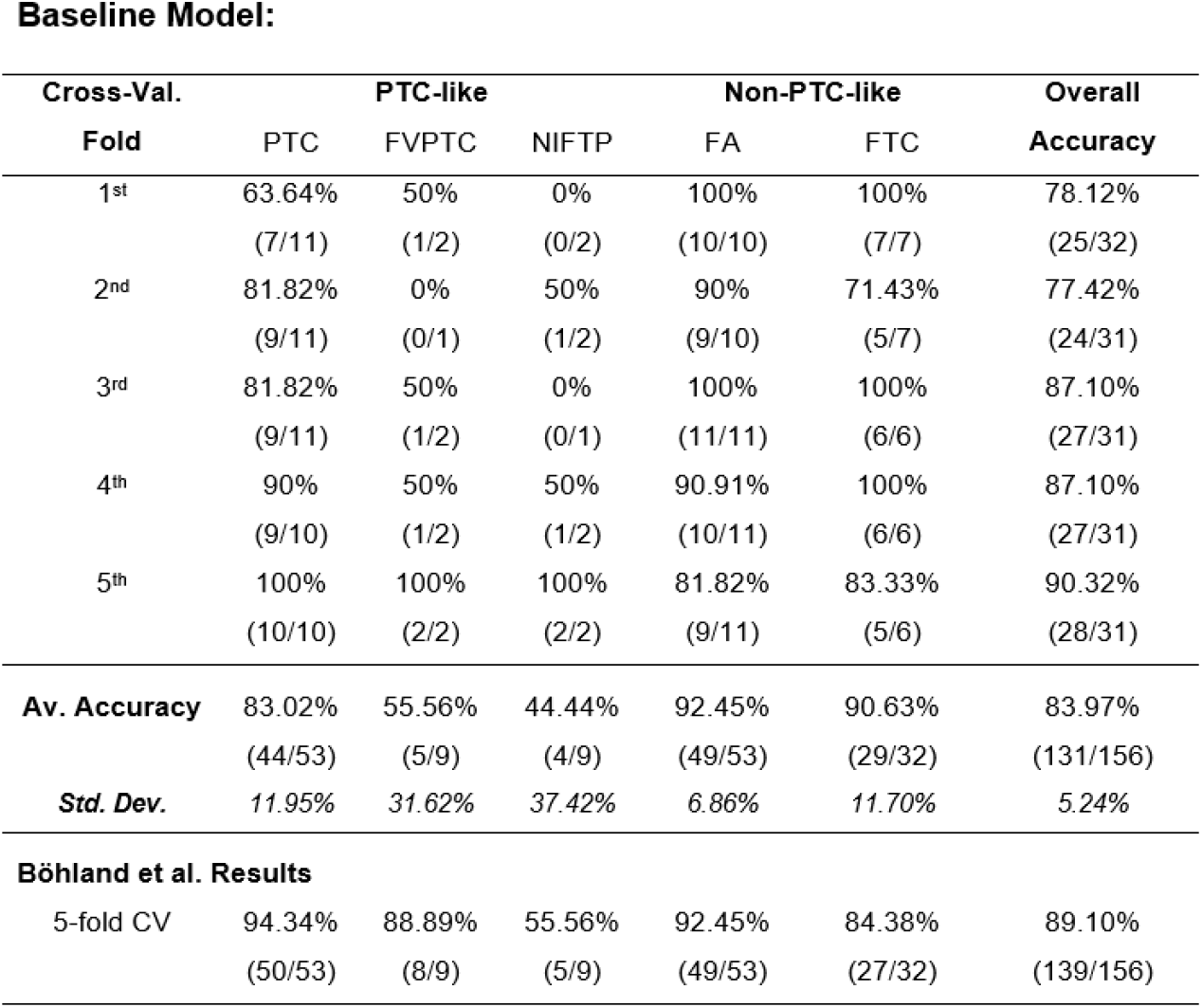
Full cross-validated results for each fold when the baseline model was tested on the T&T dataset.

**Supplementary Material, Figure 7.**
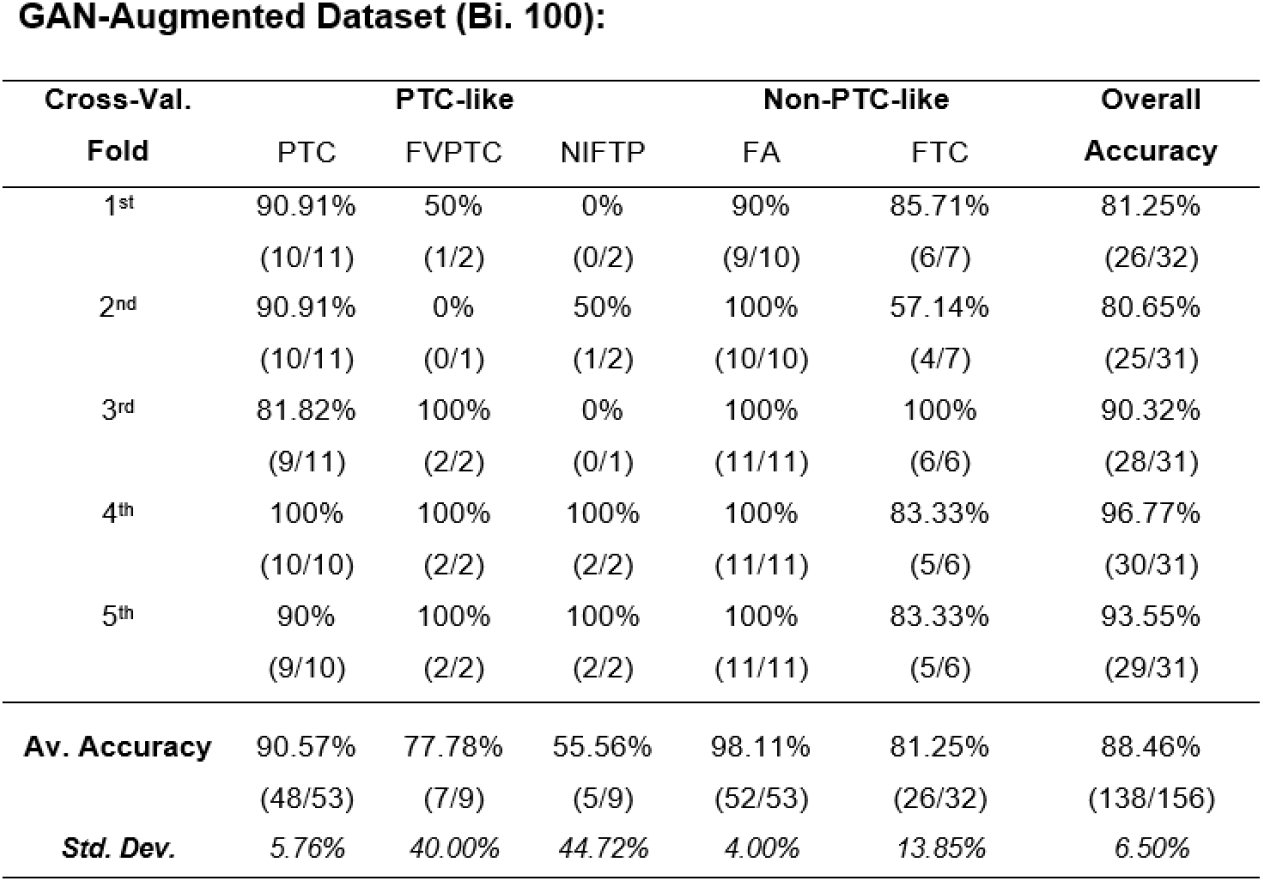
Full cross-validated results for each fold when the Binary 100 GAN-augmented model was tested on the T&T dataset.

**Supplementary Material, Figure 8.**
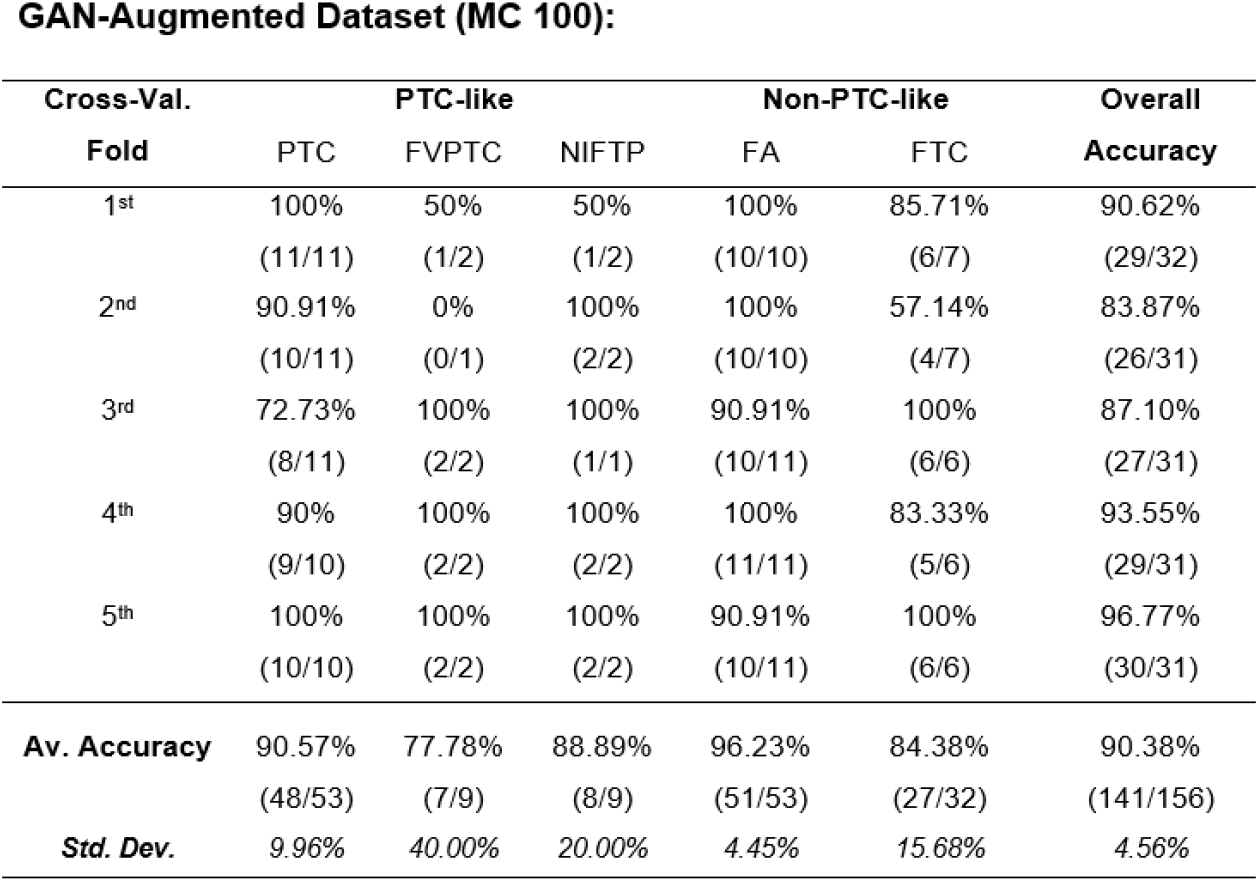
Full cross-validated results for each fold when the MC 100 GAN-augmented model was tested on the T&T dataset.

### S7. StyleGAN2-ADA Architectures

Summaries of the StyleGAN2-ADA pytorch generator and discriminator model architectures are included within this section.

**Supplementary Material, Figure 9.**
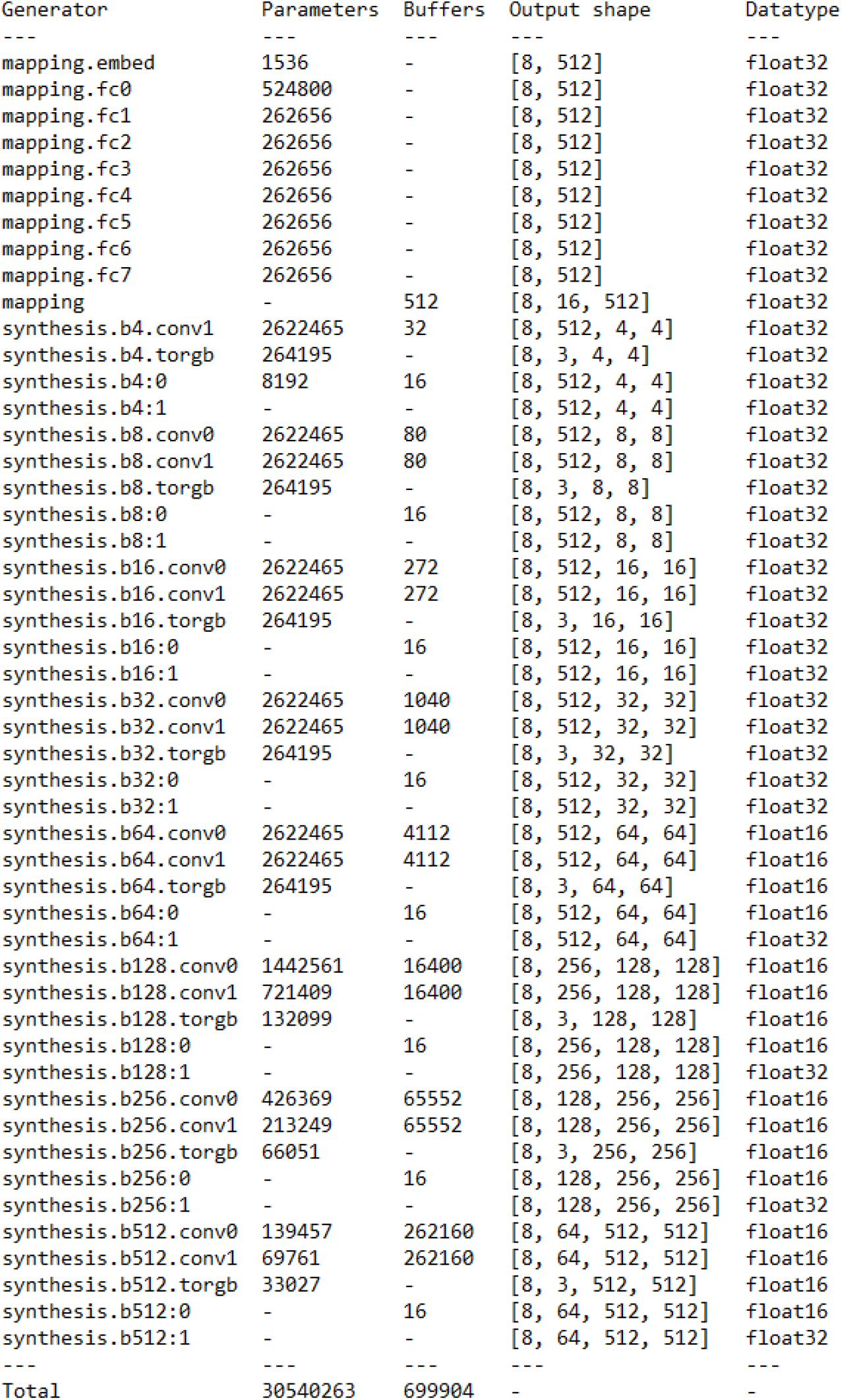
A summary of the StyleGAN2-ADA pytorch generator architecture.

**Supplementary Material, Figure 10.**
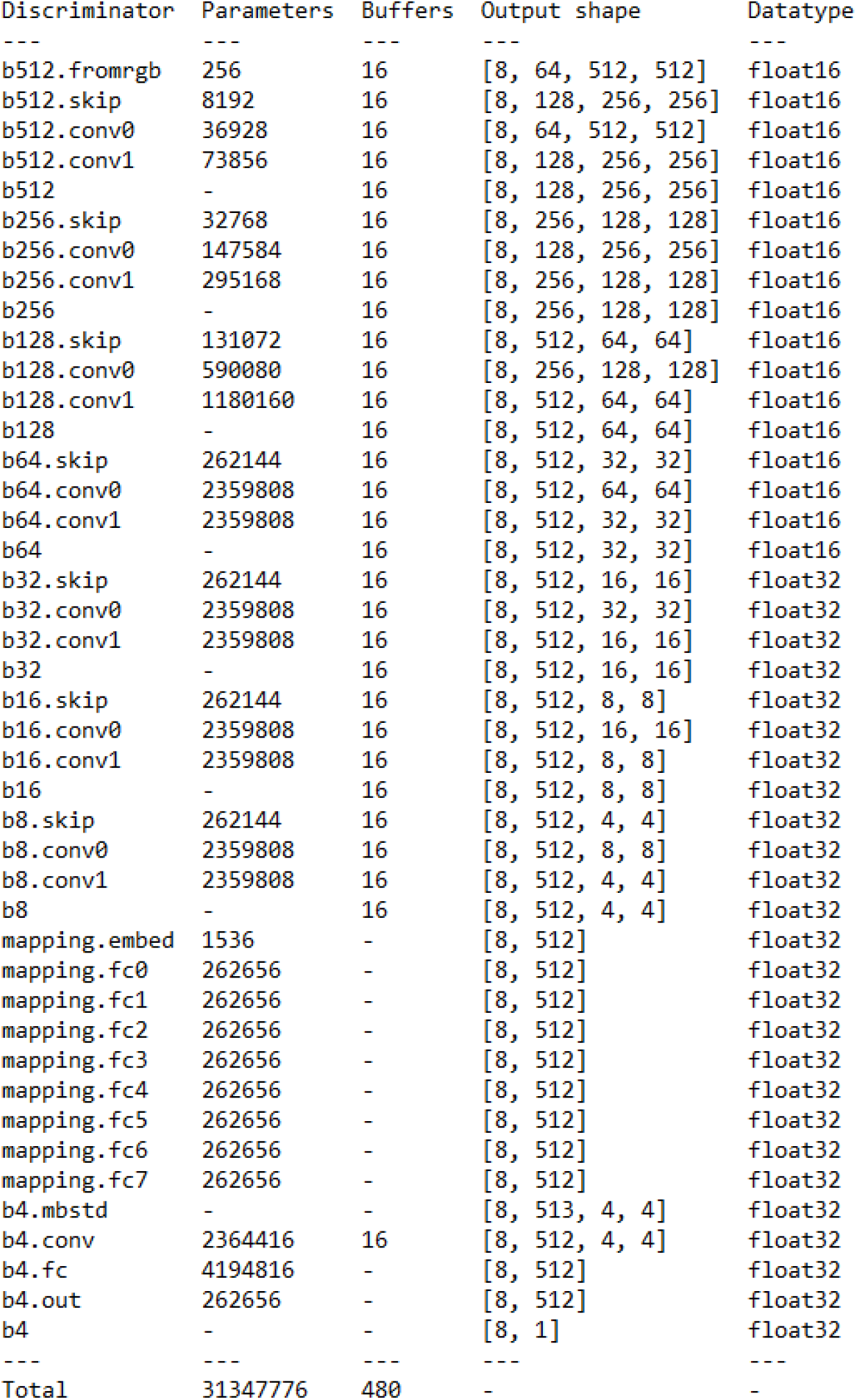
A summary of the StyleGAN2-ADA pytorch discriminator architecture.

## Notes

https://github.com/williamdee1/ThyCa-GAN

### Competing Interest Statement

The authors have declared no competing interest.

